# Cooperative Stabilization of Close-Contact Zones Leads to Sensitivity and Selectivity in T-Cell Recognition

**DOI:** 10.1101/2021.03.02.433515

**Authors:** Bartosz Różycki, Thomas R. Weikl

## Abstract

T cells are sensitive to 1 to 10 foreign-peptide-MHC complexes among a vast majority of self-peptide-MHC complexes, and discriminate selectively between peptide-MHC complexes that differ not much in their binding affinity to T-cell receptors (TCRs). Quantitative models that aim to explain this sensitivity and selectivity largely focus on single TCR/peptide-MHC complexes, but T cell adhesion involves a multitude of different complexes. In this article, we demonstrate in a three-dimensional computational model of T-cell adhesion that the cooperative stabilization of close-contact zones is sensitive to 1 to 3 foreign-peptide-MHC complexes and occurs at a rather sharp threshold affinity of these complexes, which implies selectivity. In these close-contact zones with lateral extensions of hundred to several hundred nanometers, few TCR/foreign-peptide-MHC complexes and many TCR/self-peptide-MHC complexes are segregated from LFA-1/ICAM-1 complexes that form at larger membrane separations. Previous high-resolution microscopy experiments indicate that the sensitivity and selectivity in the formation of closed-contact zones reported here is relevant for T-cell recognition, because the stabilization of close-contact zones by foreign, agonist peptide-MHC complexes precedes T-cell signaling and activation in the experiments.

## 1. Introduction

T cells recognize foreign peptides that are bound to major histocompatibility complexes (MHCs) on apposing cell surfaces among a vast majority of self peptides [1]. This recognition is highly sensitive and selective: T cells can be sensitive to between 1 and 10 foreign-peptide-MHC complexes [2–5], and discriminate selectively between foreign-peptide-MHC and self-peptide-MHC complexes that differ not strongly in their binding affinity to T-cell receptors [6–9]. Such selectivity is typically seen to require kinetic proof-reading in TCR binding, which involves a series of biochemical transformations in TCR/peptide-MHC complexes [10–13]. Kinetic proofreading focuses on individual TCR binding events and single TCR/peptide-MHC complexes, but T cells interact with apposing cells via many complexes. Increasing the concentration of self-peptide-MHCs has been shown to increase the response of T cells to foreign-peptide-MHCs [14,15], which points to a cooperativity between TCR/self-peptide-MHC and TCR/foreign-peptide-MHC complexes in T-cell recognition. The formation of these TCR complexes occurs at close contact of about 15 nm, while the longer LFA-1/ICAM-1 complexes form at larger cell-cell separations of about 40 nm [16]. High-resolution microscopy experiments showed that TCR/foreign-peptide-MHC complexes stabilize close-contact zones during the scanning of antigen-presenting cells by T cells [17]. The stabilization of the close-contact zones is independent of the actin cytoskeleton and of TCR-induced signaling and, thus, appears to precede and initiate T-cell signaling and activation.

In this article, we investigate the interplay of few TCR/foreign-peptide-MHC and many TCR/self-peptide-MHC and LFA-1/ICAM-1 complexes in a computational model of T-cell adhesion. The longer LFA-1/ICAM-1 complexes tend to segregate from the shorter TCR/MHC complexes because the membranes need to curve to compensate the length mismatch, which costs bending energy. The TCR/foreign-peptide-MHC and TCR/self-peptide-MHC complexes therefore form in close-contact zones that are separated from domains of LFA-1/ICAM-1 complexes. Figure 1A illustrates adhesion in the absence of foreign-peptide-MHC complexes and at a binding energy of TCR/self-peptide-MHC complexes at which close-contact zones are not stable. The adhesion is then mediated only by LFA-1/ICAM-1 complexes (grey dots). In Figure 1B, the addition of just three foreign-peptide-MHC complexes with larger binding energy to TCRs leads to a close-contact zone that is cooperatively stabilized by few TCR/foreign-peptide-MHC complexes (black dots) and many TCR/self-peptide-MHC complexes (light grey dots). We systematically vary the binding energies of the TCR/self-peptide-MHC and TCR/foreign-peptide-MHC complexes and find that the cooperative stabilization of closed contact zones occurs selectively at a rather sharp threshold value of the TCR/foreign-peptide-MHC binding energy.

**Figure 1.**
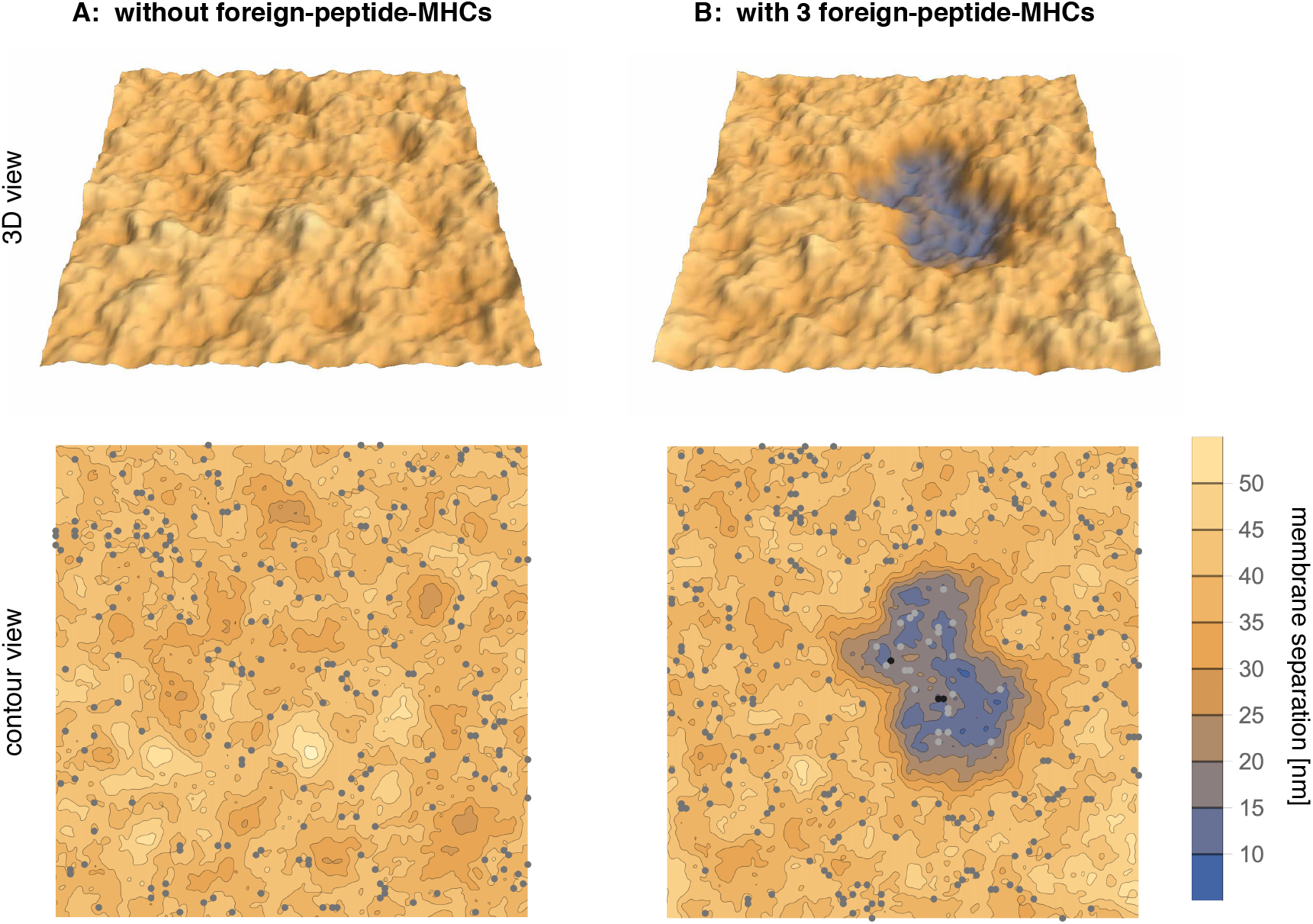
Simulation conformations of adhering membrane segments with area 1.5 *×* 1.5 *μ*m^2^ (A) in the absence of foreign-peptide-MHCs and (B) in the presence of 3 foreign-peptide MHCs with binding energy *U*_*f*_ = 12 *k*_*B*_*T* to TCRs. In (A), the binding energy *U*_*s*_ = 5.9 *k*_*B*_*T* of the 270 self-peptide-MHCs is not sufficient to stabilize a close-contact zone of the adhering membranes. The adhesion is therefore mediated by LFA-1/ICAM-1 complexes (grey dots) with a concentration of about 100/*μ*m^2^. The close-contact zone in (B) is cooperatively stabilized by 3 TCR/foreign-petide-MHC complexes (black dots) and many TCR/self-peptide-MHC complexes (light grey dots).

## 2. Results

We first consider an adhesion scenario in which TCRs can only bind to one type of peptide-MHC complexes. This scenario corresponds to T-cell adhesion on supported membranes that contain one type of peptide-MHCs and ICAM-1 [17–19]. In our model, the two adhering membranes are discretized into apposing patches that can contain single proteins. The proteins diffuse along the membranes by hopping from patch to patch, and bind to proteins at apposing membrane patches if the separation of the patches is within 15 ± 0.5 nm for TCR/MHC complexes and within 40 ± 0.5 nm for LFA-1/ICAM-1 complexes. The membrane bending energy associated with variations in the membrane separation depends on the effective bending rigidity *κ* = *κ*_1_*κ*_2_/(*κ*_1_ + *κ*_2_) of the two membranes with rigidities *κ*_1_ and *κ*_2_ [20,21]. We use the value *κ* = 20 *k*_*B*_*T* in accordance with typical values of bending rigidities of lipid membranes between 10 and 40 *k*_*B*_*T* [22,23]. Our membranes consist of 100 × 100 patches with a projected area of 15 × 15 nm^2^ and, thus, have a total projected area of 1.5 × 1.5 *μ*m^2^. In the adhesion scenario with one type of peptide-MHC complexes, the T-cell membrane contains 270 TCR and 270 LFA-1 proteins, and the apposing membrane contains 270 peptide-MHC and 270 ICAM-1 proteins. The total concentration of all four protein species therefore is 120/*μ*m^2^. We adjust the binding energy of LFA-1/ICAM-1 complexes to 9.5 *k*_*B*_*T*, which leads to a concentration of the complexes of about 100/*μ*m^2^ [18] for small numbers of TCR/peptide-MHC complexes, and systematically vary the binding energy *U* of the TCR/peptide-MHC complexes. For each value of *U*, we run 6 Monte Carlo (MC) simulations with a length of 2 · 10^8^ *t*_*o*_ where *t*_*o*_ is the time associated with an MC step. In an MC step, we attempt to shift the separation of each pair of apposing membrane patches and to translate each protein and each pair of apposing binding partners to a nearest-neighbor patch along the membrane in independent substeps. To sample the equilibrium adhesion behavior, we discard the initial 4 · 10^7^ MC steps of the MC simulations during which the adhering membranes relax into equilibrium from different initial conformations (see Methods for details).

Figure 2 A shows how the average number of TCR/peptide-MHC complexes depends on the binding energy *U* of the complexes. The number of TCR/peptide-MHC complexes increases strongly at a binding energy *U* of about 6.0 *k*_*B*_*T* at which a close-contact zone is beginning to be stabilized by the TCR/peptide-MHC complexes. At binding energies *U* ≥ 6.5 *k*_*B*_*T*, the probability for such a close-contact zone is 1 (see Figure 2B), which means that a close-contact zone of TCR/peptide-MHC complexes is continuously present in the equilibrated simulations. The lifetime of the close-contact zone therefore becomes equal to the length *T*_*o*_ = 1.6 · 10^8^ *t*_*o*_ of the equilibrated trajectory parts over which these quantities are measured (see Figure 2D). The probability and lifetime of close-contact zones is calculated from simulation conformations at intervals of 2 *·* 10^5^*t*_*o*_ in the equilibrated trajectory parts. We define a close-contact event as a contiguous sequence of conformations (i) that contains pairs of apposing membrane patches with separation less than 20 nm in each conformation and (ii) in which TCR/peptide-MHC complexes are present in at least one conformation in the sequence. The probability of close-contact zones is the fraction of conformations that are part of close-contact events. This definition of close-contact zones takes into account that the instantaneous number of TCR/peptide-MHC complexes in a small close-contact zone with few complexes can briefly drop to 0 because of fluctuations in the number of complexes from binding and unbinding events. The average area of the close-contacts zones grows for binding energies *U* ≥ 6.0 *k*_*B*_*T* at which the number of TCR/peptide-MHC increases (see Figure 2C).

**Figure 2.**
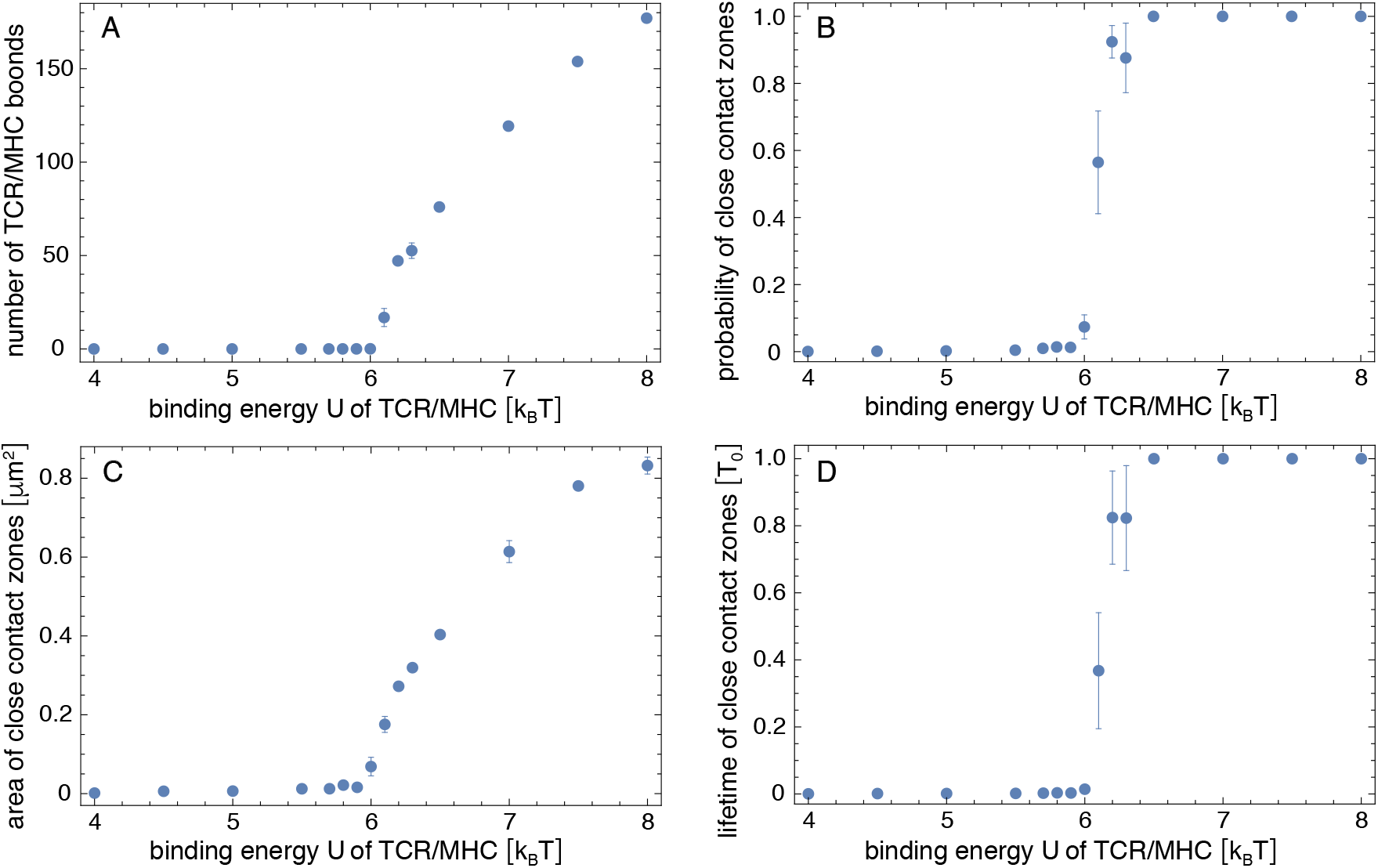
Simulation results in the adhesion scenario with only one type of peptide-MHC complexes. The (A) number of TCR/peptide-MHCs complexes, and the (B) probability, (C) area, and (D) lifetime of close-contact zones are average values obtained from the equilibrated simulation trajectories of length *T*_*o*_ = 1.6 *·* 10^8^ *t*_*o*_ where *t*_*o*_ is the time corresponding to one MC simulation step. The errors are calculated as the error of the mean for 6 independent trajectories at each value of the binding energy *U* of the TCR/peptide-MHC complexes.

We now consider adhesion scenarios in which TCRs can bind to two types of peptide-MHCs: to self-peptide-MHCs with binding energy *U*_*s*_, or to foreign-peptide-MHCs with larger binding energy *U*_*f*_. This situation corresponds to T-cell adhesion on supported membranes that contain self-peptide-MHCs, foreign-peptide-MHCs, and ICAM-1 [14], and mimics the interplay of self- and foreign-peptide-MHC complexes in the adhesion of T cells to antigen-presenting cells. In our simulation systems, we either have 3 foreign-peptide-MHCs and 267 self-peptide-MHCs, or 1 foreign-peptide-MHC and 269 self-peptide-MHCs. The concentration of foreign-peptide-MHCs thus is 1.3/*μ*m^2^ or 0.4/*μ*m^2^, respectively, for the membrane area 1.5 × 1.5 *μ*m^2^ of our simulations. As before, the numbers of TCRs, LFA-1 proteins, and ICAM-1 proteins are 270 each, and the binding energy of LFA-1/ICAM-1 complexes is 9.5 *k*_*B*_*T*. We assume that the self-peptide-MHCs alone to do not stabilize a close-contact zone and consider the three values *U*_*s*_ = 4.0, 5.5, and 5.9 *k*_*B*_*T* for the binding energy of TCR/self-peptide-MHC, which are below the threshold value of about 6.0 *k*_*B*_*T* for stable contact zones with one type of peptide-MHCs (see Figure 2).

Figure 3 shows how the probability and lifetime of close-contact zones depends on the binding energy *U*_*f*_ of the TCR/foreign-peptide-MHC complexes. In our simulations with 3 foreign-peptide-MHCs, the lifetime of close-contact zones increases rather strongly at threshold values of *U*_*f*_ that depend on the binding energy *U*_*s*_ of the TCR/self-peptide-MHC complexes. For *U*_*s*_ = 4.0 *k*_*B*_*T*, the lifetime of the close-contacts zones increases from 0.023 *±* 0.005 *T*_*o*_ at *U*_*f*_ = 12 *k*_*B*_*T* to 0.30 *±* 0.14 *T*_*o*_ at *U*_*f*_ = 13 *k*_*B*_*T* and, thus, by a factor of 13 *±* 7 with an increase of 1 *k*_*B*_*T* in *U*_*f*_. For *U*_*s*_ = 5.5 *k*_*B*_*T*, the lifetime increases by a factor of 10 *±* 3 from 0.012 *±* 0.001 *T*_*o*_ at *U*_*f*_ = 11 *k*_*B*_*T* to 0.11 *±* 0.03 *T*_*o*_ at *U*_*f*_ = 12 *k*_*B*_*T*. And for *U*_*s*_ = 5.9 *k*_*B*_*T*, the lifetime increases by a factor of 8 *±* 5 from 0.06 *±* 0.03 *T*_*o*_ at *U*_*f*_ = 10 *k*_*B*_*T* to 0.47 *±* 0.15 *T*_*o*_ at *U*_*f*_ = 11 *k*_*B*_*T*. These increases in the lifetimes of close-contact zones by a factor of about 10 for an increase of 1 *k*_*B*_*T* in the binding energy *U*_*f*_ for the 3 foreign-peptide-MHCs are significantly larger than the increases in the lifetimes of the individual complexes. In our simulations, the lifetimes of TCR/peptide-MHC complexes is proportional to exp[*U*/*k*_*B*_*T*], irrespective of the stability of the close-contact zones in which these complexes form (see Figure 4B), and thus increase by a factor of 2.7 with an increase of 1 *k*_*B*_*T* in the binding energy *U*. For large values of *U*_*f*_, the close-contact zones contain 3 TCR/foreign-peptide-MHC complexes and on average about 21 TCR/self-peptide-MHC complexes for *U*_*s*_ = 5.9 *k*_*B*_*T*, about 5 TCR/self-peptide-MHC complexes for *U*_*s*_ = 5.5 *k*_*B*_*T*, and on average about 0.5 TCR/self-peptide-MHC complexes for *U*_*s*_ = 4.0 *k*_*B*_*T* (see Figures 4A, 1B, and 5A,B). The close-contact zones thus are jointly stabilized by TCR/foreign-peptide-MHC and TCR/self-peptide-MHC complexes for *U*_*s*_ = 5.9 *k*_*B*_*T* and 5.5 *k*_*B*_*T*, and predominantly by the TCR/foreign-peptide-MHC complexes for *U*_*s*_ = 4.0 *k*_*B*_*T*. The threshold value of *U*_*f*_ at which the lifetime of the close-contact zones increases rather strongly is reduced by about 2.5 to 3 *k*_*B*_*T* for an increase of *U*_*s*_ from 4.0 to 5.9 *k*_*B*_*T* (see Figure 3B).

**Figure 3.**
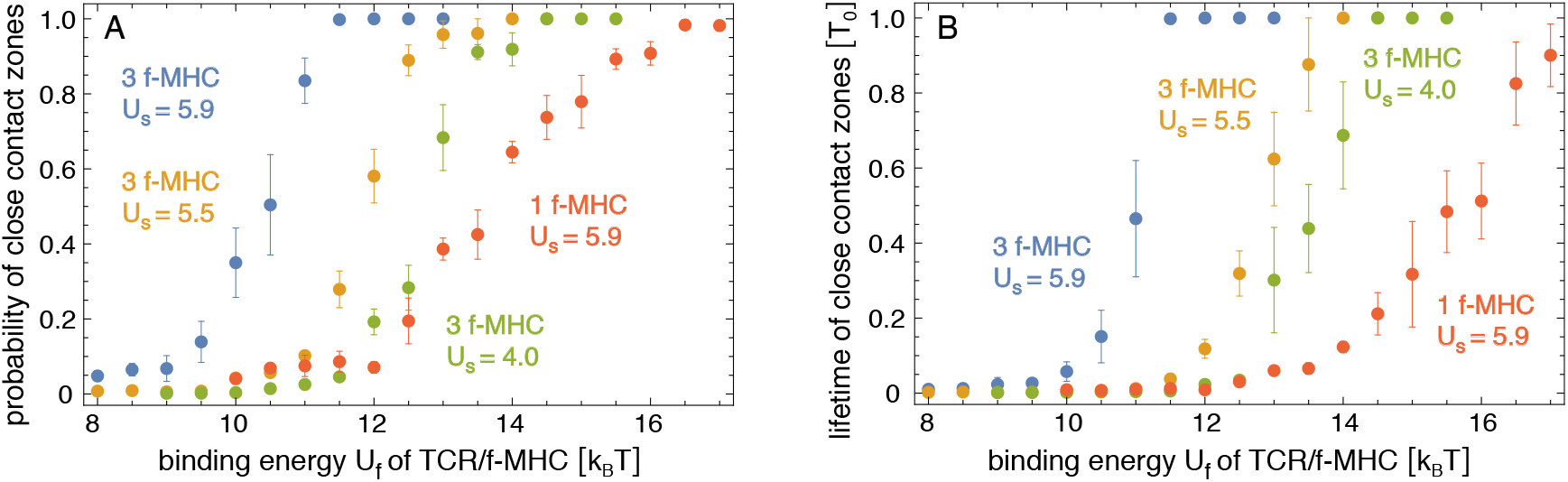
(A) Probability and (B) lifetime of close-contact zones in adhesion scenarios with either 3 or 1 foreign-peptide-MHC (f-MHC) *versus* binding energy *U*_*f*_ of TCR/f-MHC complexes, for different values of the binding energy *U*_*s*_ of TCR/self-peptide-MHCs in units of the thermal energy *k*_*B*_*T*. The errors are calculated as the error of the mean for 6 independent trajectories at each value of *U*_*f*_ in the different simulation systems.

**Figure 4.**
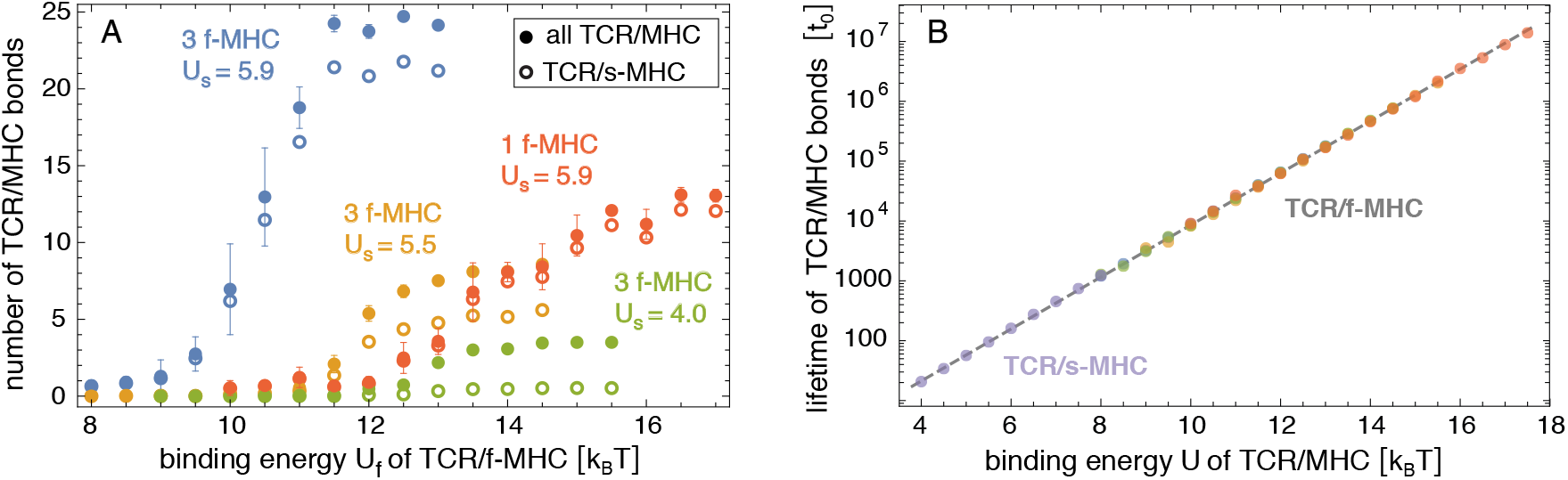
(A) Average number of all TCR/peptide-MHC and of TCR/self-peptide-MHC complexes *versus* binding energy *U*_*f*_ of TCR/foreign-peptide-MHC complexes in adhesion scenarios with either 3 or 1 foreign-peptide-MHC (f-MHC)and different values of the binding energy *U*_*s*_ of TCR/self-peptide-MHCs in units of the thermal energy *k*_*B*_*T*. (B) Average lifetime of TCR/peptide-MHC complexes *versus* binding energy of the complexes in all adhesion scenarios. The lifetimes of the TCR/self-peptide-MHCs in the adhesion scenario without foreign-peptide-MHCs are shown in purple. The lifetimes of the TCR/foreign-peptide-MHCs in the adhesion scenarios with either 3 or 1 foreign-peptide-MHC are shown in the same colors as in (A). The dashed line is the regression line exp(*c* + *U*/*k*_*B*_*T*)*t*_*o*_ of all data points with the single fit parameter *c* = *−*0.945 *±* 0.007.

**Figure 5.**
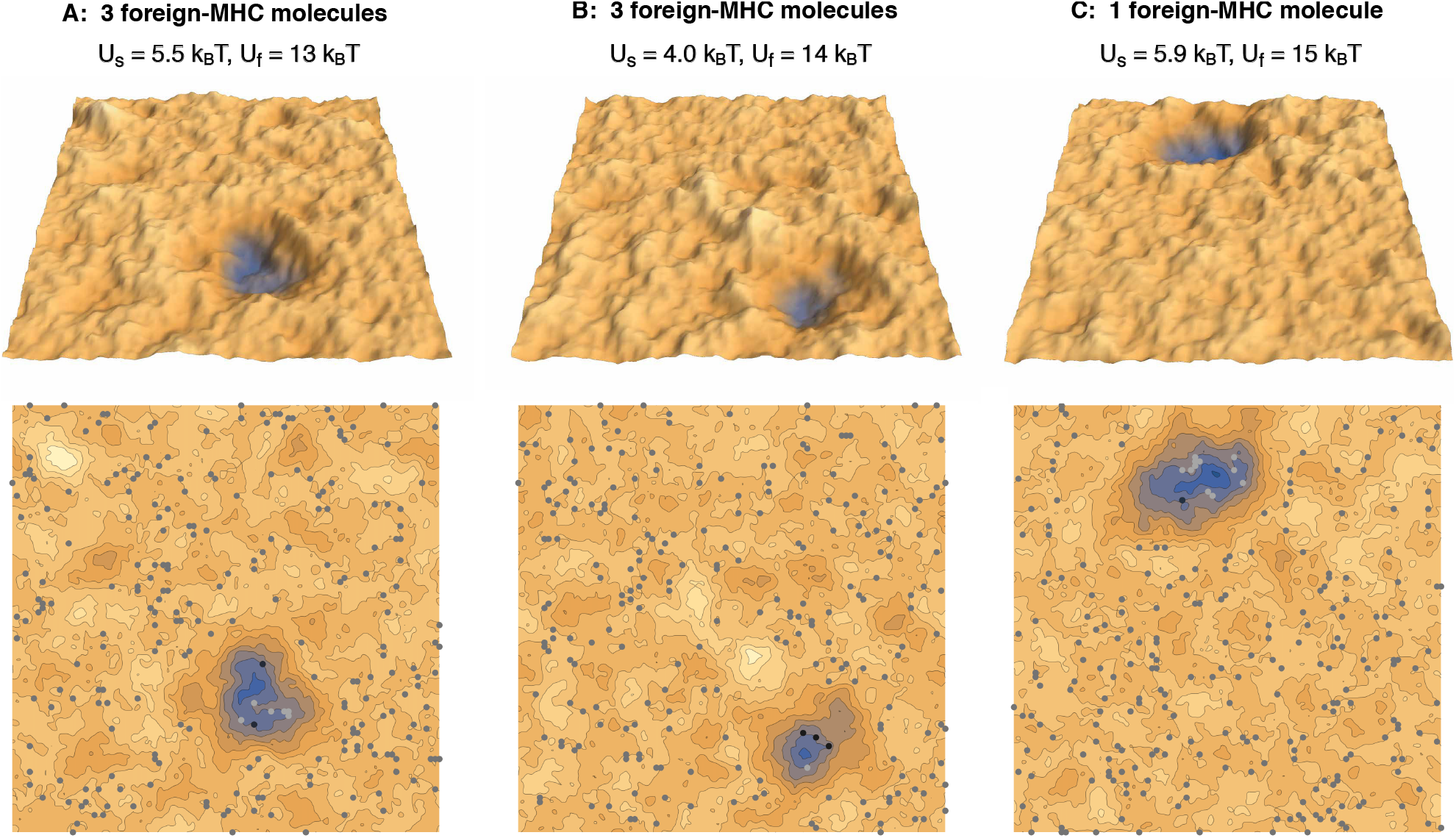
Simulation conformations of membrane segments with area 1.5 *×* 1.5 *μ*m^2^ in adhesion scenarios with (A,B) three foreign-peptide-MHCs and (C) a single foreign-peptide-MHC. The different complexes and different values of the membrane separation are indicated in the same colors as in Figure 1.

In our simulations with 1 foreign-peptide-MHC complex, the lifetime of close-contact zones increases by about a factor of 3 with an increase of 1 *k*_*B*_*T* in *U*_*f*_ in the range from *U*_*f*_ = 12 *k*_*B*_*T* to 15 *k*_*B*_*T*, which is comparable to the increase in the lifetime of the TCR/foreign-peptide-MHC complex with *U*_*f*_. However, the lifetimes of the close-contact zones is about a factor 40 larger than the lifetime of the TCR/foreign-peptide-MHC complex in this range of *U*_*f*_ values, which reflects the cooperative stabilization of close-contact zones by the TCR/self-peptide-MHC complexes with binding energy *U*_*s*_ = 5.9 *k*_*B*_*T* in our simulations. For large values *U*_*f*_ ≥ 15 *k*_*B*_*T*, the close-contact zones contain between 10 and 12 TCR/self-peptide-MHC complexes, besides the single TCR/foreign-peptide-MHC complex in these simulations (see Figures 4A and 5C).

## 3. Discussion

Our simulation results indicate that the cooperative stabilization of close-contact zones is sensitive to few TCR/foreign-peptide-MHC complexes. In our simulations with 3 foreign-peptide-MHCs, a clear stabilization of close-contact zones occurs within a rather narrow window of about 2 *k*_*B*_*T* in the binding energy *U*_*f*_ of the TCR/foreign-peptide-MHC complexes, which implies selectivity between TCR/peptide-MHC complexes at both sides of these windows and, thus, between peptide-MHC complexes that differ by only 2 *k*_*B*_*T* in their binding energies. In the adhesion scenario with 3 foreign-peptide-MHCs and the binding energy *U*_*s*_ = 5.5 *k*_*B*_*T* of TCR/self-peptide-MHC complexes, the probability and lifetime of close-contact zones changes significantly in the 2 *k*_*B*_*T* window from about *U*_*f*_ = 10.5 *k*_*B*_*T* to 12.5 *k*_*B*_*T*. If we assume that a binding energy *U*_*f*_ = 12.5 *k*_*B*_*T* corresponds to a TCR/foreign-MHC-peptide complex with a typical lifetime of 1 s [19,24], the time step *t*_*o*_ of our MC simulations corresponds to a physical time of 1 *μ*s according to the regression line in Figure 4B. The average lifetime of close contact zones of about 0.3 *T*_*o*_ at *U*_*f*_ = 12.5 *k*_*B*_*T* then corresponds to a physical time of 50 s. In contrast, the average lifetime of close contact zones at *U*_*f*_ = 10.5 *k*_*B*_*T* is then only 1 s. In addition, the average probability of close-contacts zone at *U*_*f*_ = 10.5 *k*_*B*_*T* is about 6% and, thus, signicantly smaller than the probability of 0.89% at *U*_*f*_ = 12.5 *k*_*B*_*T*. These changes in the probability and lifetime of close contact zones are large compared to the change by a factor of about 7 in the lifetime of individual TCR/MHC-peptide complexes for an increase of 2*k*_*B*_*T* in the binding energy. In the adhesion scenario with 1 foreign-peptide-MHC, a comparable increase in the probability of the close contact zones occurs within a wider window of 4 *k*_*B*_*T* from about *U*_*f*_ = 11 *k*_*B*_*T* to *U*_*f*_ = 15 *k*_*B*_*T* (see Figure 3A).

In our adhesion scenario with 3 foreign-peptide-MHCs, the window of *U*_*f*_ values in which the stabilization of the close-contact zones occurs depends on the binding energy *U*_*s*_ of TCR/self-peptide-MHC complexes. At the binding energy *U*_*s*_ = 4.0 *k*_*B*_*T*, close contact zones are predominantly stabilized by TCR/foreign-peptide-MHC complexes alone. TCR/self-peptide-MHC complexes only form with low probability and, thus, do not contribute significantly to the stabilization. At the binding energies *U*_*s*_ = 5.5 *k*_*B*_*T* and *U*_*s*_ = 5.9 *k*_*B*_*T*, the close-contact zones are jointly stabilized by TCR/foreign-peptide-MHC and TCR/self-peptide-MHC complexes, and the stabilization window is shifted to smaller values of *U*_*f*_. The joint stabilization of close contact zones by TCR/foreign-peptide-MHC and TCR/self-peptide-MHC complexes is in line with experimental findings that self-peptide-MHCs increase and facilitate the response of T cells to foreign-peptide-MHCs [14,15]. The cooperativity of self-peptide-MHCs and foreign-peptide-MHCs has been suggested to result from positive selection of naive T cells with TCRs that interact with self-peptide-MHCs on antigen-presenting cells in the thymus [15]. This positive selection is balanced by negative selection to ensure that T cells are not activated by self-peptide-MHCs alone [25].

The membrane separation in closed-contact zones varies between about 10 nm and 20 nm in our simulations (see Figures 1B and 5). Proteins with extracellular protrusions larger than 25 nm such as CD45 therefore are rather clearly excluded from these close-contact zones as suggested in the kinetic-segregation model of T-cell activation. In this model, the size-based segregation of the inhibitory tyrosine phosphatase CD45 from TCR complexes in close-contact zones triggers T-cell signaling and activation [26–29]. Our simulation results can also be seen to connect to the serial-engagement model that suggests that foreign-peptide-MHC complexes bind to many different TCRs [30,31]. The lifetimes of close-contact zones in our simulations are much larger then the lifetime of the individual TCR/foreign-peptide-MHC complexes that stabilize the close-contact zones, which implies many binding and unbinding events of foreign-peptide-MHC complexes during the lifetime of a close contact-zone.

While the lifetimes of TCR/peptide-MHC complexes depends only on the binding energy *U* in our simulations, the two-dimensional binding constant *K*_2D_ is also affected by the distribution *P*(*l*) of local membrane separations. The two-dimensional binding constant can be calculated as *K*_2D_ = *∫ k*_2D_(*l*)*P*(*l*)d*l* where *k*_2D_(*l*) is the binding constant as a function of the separation *l* [32]. In our model, *k*_2D_(*l*) is equal to *a*^2^ exp[*U*/*k*_*B*_*T*] for separations *l* within the binding range 15 ± 0.5 nm of TCR/peptide-MHC complexes, where *a*^2^ = 15 × 15 nm^2^ is the area of a membrane patch, and is equal to 0 for separations *l* outside this binding range [33]. In large close-contact zones of our simulations, about 20% of the membrane patches are within the binding range of TCR/peptide-MHC complexes. The two-dimensional binding constant of these complexes therefore is *K*_2D_ = *k*_2D_(*l*)*P*(*l*)d*l* ≃ 0.2 *a*^2^ exp[*U*/*k*_*B*_*T*], which is about 10 *μ*m^2^ for the binding energy *U* = 12.5 *k*_*B*_*T* of the example above. Within the domains of LFA-1/ICAM-1 complexes, the binding constant *K*_2D_ for TCR/peptide-MHC complexes is 0 simply because the distribution *P*(*l*) of local membrane separations within these domains does not allow for the binding of TCR/peptide-MHC complexes.

TCR/peptide-MHC and LFA-1/ICAM-1 complexes tend to segregate in our model because the membranes need to curve to compensate the length mismatch [34]. Experiments in the last years highlight the importance of size and length in the segregation of complexes and proteins in membrane adhesion [35–39]. In particular the clustering of the initially randomly distributed TCRs [40] during T-cell adhesion has been a focus in understanding T-cell activation [41–46] Our previous simulations and calculations indicate that the curvature-mediated segregation into domains of LFA-1/ICAM-1 complexes and close-contact zones of TCR/peptide-MHC complexes occurs for concentrations

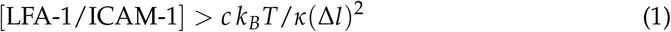

of LFA-1/ICAM-1 complexes with length difference Δ*l* to TCR/peptide-MHC complexes and the numerical prefactor *c* = 0.65 ± 0.15 [20,47,48]. For *κ* = 20 *k*_*B*_*T* and Δ*l* = 25 nm as in our model, the critical concentration of segregation at the right-hand side of this inequality is about 50/*μ*m^2^. The concentration [LFA-1/ICAM-1] ≃ 100/*μ*m^2^ in our model is clearly larger than this critical concentration, in agreement with the concentrations of LFA-1/ICAM-1 complexes measured in the immunological synapse of T cells on supported membranes [18]. In the adhesion of T cells to antigen-presenting cells, other complexes with comparable length to TCR/peptide-MHC complexes such as CD2 complexes and complexes between the co-receptors CD4 or CD8 with MHC may contribute to the segregation and the stabilization of close-contact zones. CD2 has been reported to enhance the response of T cells to antigens [49]. At sufficiently high concentrations, CD2 complexes more recently have been found to form peripheral domains in the immunological synapse of T cells on supported membranes [50].

High-resolution microscopy of T-cell adhesion shows that close-contact zones form at the tips of microvilli that protrude from T cells during the scanning of antigen-presenting cells [17]. These tips have a width of about 500 nm, which is larger or comparable to the width of the close-contact zones in our simulations. Close-contact zones at the microvilli tips stabilized by TCR/foreign-peptide-MHC remain after the drug-induced disassembly of the actin cytoskeleton of the T cells, which indicates that the stabilization of the close-contact zones observed in the experiments is independent of cytoskeletal forces. With intact cytoskeleton, T cells exert forces that likely play a role in bringing the cell surfaces to distances of about 40 nm, at which the LFA-1/ICAM-1 can form, against the repulsion of glycocalyx components longer than 40 nm [16]. Our simulations indicate that the nucleation of close-contact zones within the domains of LFA-1/ICAM-1 is rather fast and, thus, does not require force. However, transversal forces on TCR/foreign-peptide-MHC complexes observed in recent experiments [51], which are presumably induced by lateral motion of microvilli, may contribute to T-cell signaling and activation [24,52–56].

## 4. Methods

In our computational model of T-cell adhesion, the two adhering membrane segments are discretized into 100 × 100 apposing pairs of membrane patches [57]. The patches at opposing boundaries of the membrane segments are connected by periodic boundary conditions. Each membrane patch can contain a single protein. Our MC simulations consist of MC moves in which we attempt (1) to shift the separation of a pair of apposing membrane patches, (2) to translate a single protein to a nearest-neighbor patch, and (3) to translate a pair of apposing partner proteins to a nearest-neighbor pair of apposing patches. In the MC moves (1), a pair *i* of apposing patches is selected randomly, and the local separation *l*_*i*_ of the patches is attempted to be shifted to *l*_*i*_ + *δl* where *δl* is a random length that is distributed uniformly between 0.5 nm and 0.5 nm. In the MC moves (2), a single protein is randomly selected and attempted to be shifted to one of the four nearest-neighbor patches in the discretized membranes, provided this patch is not occupied by another protein. In the MC moves (3), a pair of apposing partner proteins, i.e. a pair of LFA-1 and ICAM-1 proteins or a pair of TCR and peptide-MHC proteins located in two apposing membrane patches, is selected randomly and independently of the separation of the patches, and is attempted to be shifted to a nearest-neighbor pair of apposing, unoccupied membrane patches. All three types MC moves can lead to the binding or unbinding of protein complexes. The MC moves (2) lead to the diffusion of single proteins, and the MC moves (3) to the diffusion of bound protein complexes. To correctly capture the lifetimes of bound complexes, unrealistic MC moves of type (2) in which a bound protein would directly hop into a new complex at a neighboring site are excluded.

The MC moves are accepted or rejected with probabilities that depend on the energy change Δ*E* associated with the move. We use the standard Metropolis criterion in which MC moves are accepted with probability 1 for Δ*E <* 0, i.e. if the moves decrease the overall energy, and with probability exp[−Δ*E*/*k*_*B*_*T*] for Δ*E >* 0. The energy change of MC moves of type (1) associated with changes in the bending energy of the membrane is calculated from the discretized effective bending energy

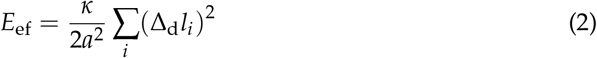

of the two membranes 1 and 2 with effective bending rigidity *κ* = *κ*_1_*κ*_2_/(*κ*_1_ + *κ*_2_) and the discretized Laplacian Δ_*d*_*l*_*i*_ = *l*_*i*1_ + *l*_*i*2_ + *l*_*i*3_ + *l*_*i*4_ *−* 4*l*_*i*_[20,21]. Here, *l*_*i*_ is the membrane separation of the apposing pair *i* of membrane patches, and *l*_*i*1_ to *l*_*i*4_ are the membrane separations at the four nearest-neighbor pairs of patches around pair *i*. In our model, the linear size *a* of the membrane patches is *a* = 15 nm as in our previous model for the cooperative binding of CD47 in the adhesion of giant plasma membrane vesicles [58]. The energy decrease and increase associated with binding and unbinding events, respectively, is simply determined from the binding energy of the complexes. In our model, an apposing pair of TCR and peptide-MHC proteins is bound if the separation *l*_*i*_ of the apposing pair *i* of membrane patches, in which the proteins are located, is within 15 *±* 0.5 nm. The binding energy of the TCR complexes is *U*_*f*_ for complexes with foreign-peptide-MHCs proteins and *U*_*s*_ for complexes with self-peptide-MHCs. An apposing pair of LFA-1 and ICAM-1 is bound with binding energy 9.5 *k*_*B*_*T* if the separation of the membrane patches is within 40 ± 0.5 nm.

A single MC step consists of 100 × 100 attempted moves of type (1), *N* attempted moves of type (2), where *N* is the total number of proteins in both membranes, and *M* attempted moves of type (3), where *M* is the instantaneous number of apposing partner proteins. A single MC trajectory consists of of 2 × 10^8^ MC steps. The dwell times of TCR-MHC bonds as well as the numbers of binding and unbinding events were computed on the fly during the second halves of the trajectories, i.e., from 10^8^ MC moves. For each set of parameter values in our adhesion scenarios, we ran six MC simulations with different initial conformations. In three of the six MC runs, the separations *l*_*i*_ of the apposing membrane patches were initially distributed randomly in the intervals from 14.5 to 40.5 nm, from 13.5 to 41.5 nm, and from 12.5 to 42.5 nm, respectively. With these initial distributions of membrane separations, both TCR/peptide-MHC complexes and the LFA-1/ICAM-1 complexes were formed at the beginning of the MC trajectories. In the three other of the six MC runs, the initial membrane separations *l*_*i*_ were distributed randomly in the intervals from 39.5 to 40.5 nm, from 38.5 to 41.5 nm, and from 37.5 to 42.5 nm, respectively. In these MC runs, the membrane adhesion was initially mediated only by the LFA-1/ICAM-1 complexes. In all MC runs, the proteins were initially distributed uniformly within the membranes. The MC simulations relax into equilibrium from these different initial conformations within the first 4 · 10^7^ MC steps of the MC simulations. We have therefore discarded these MC steps in our analysis of the equilibrium adhesion behavior.

Our three-dimensional, computational model is related to previous models that have been used to investigate the formation and temporal evolution of domain patterns in the immunological synapse of T cells [59–62]. A difference is the smaller size of the membrane patches in the simulations presented here, which allows for a higher resolution of the membrane shape and fluctuations within and between the close-contact zones of TCR/peptide-MHC complexes. In other models of the pattern formation, the proteins have been described with continuous distributions, and not as single molecules [63–66].

## References

1. Rossjohn, J.; Gras, S.; Miles, J.J.; Turner, S.J.; Godfrey, D.I.; McCluskey, J. T cell antigen receptor recognition of antigen-presenting molecules. Annu Rev Immunol 2015, 33, 169–200. doi:10.1146/annurev-immunol-032414-112334.

2. Huang, J.; Brameshuber, M.; Zeng, X.; Xie, J.; Li, Q.j.; Chien, Y.h.; Valitutti, S.; Davis, M.M. A Single Peptide-Major Histocompatibility Complex Ligand Triggers Digital Cytokine Secretion in CD4^(^+) T Cells. Immunity 2013, 39, 846–857. doi:10.1016/j.immuni.2013.08.036.

3. Sykulev, Y.; Joo, M.; Vturina, I.; Tsomides, T.J.; Eisen, H.N. Evidence that a single peptide-MHC complex on a target cell can elicit a cytolytic T cell response. Immunity 1996, 4, 565–571. doi:10.1016/s1074-7613(00)80483-5.

4. Irvine, D.J.; Purbhoo, M.A.; Krogsgaard, M.; Davis, M.M. Direct observation of ligand recognition by T cells. Nature 2002, 419, 845–849. doi:10.1038/nature01076.

5. Purbhoo, M.A.; Irvine, D.J.; Huppa, J.B.; Davis, M.M. T cell killing does not require the formation of a stable mature immunological synapse. Nat. Immunol. 2004, 5, 524–530. doi:10.1038/ni1058.

6. Krogsgaard, M.; Prado, N.; Adams, E.J.; He, X.l.; Chow, D.C.; Wilson, D.B.; Garcia, K.C.; Davis, M.M. Evidence that structural rearrangements and/or flexibility during TCR binding can contribute to T cell activation. Mol Cell 2003, 12, 1367–1378. doi:10.1016/s1097-2765(03)00474-x.

7. Daniels, M.A.; Teixeiro, E.; Gill, J.; Hausmann, B.; Roubaty, D.; Holmberg, K.; Werlen, G.; Holländer, G.A.; Gascoigne, N.R.J.; Palmer, E. Thymic selection threshold defined by compartmentalization of Ras/MAPK signalling. Nature 2006, 444, 724–729. doi:10.1038/nature05269.

8. Juang, J.; Ebert, P.J.R.; Feng, D.; Garcia, K.C.; Krogsgaard, M.; Davis, M.M. Peptide-MHC heterodimers show that thymic positive selection requires a more restricted set of self-peptides than negative selection. J. Exp. Med. 2010, 207, 1223–1234. doi:10.1084/jem.20092170.

9. Kersh, G.J.; Kersh, E.N.; Fremont, D.H.; Allen, P.M. High- and low-potency ligands with similar affinities for the TCR: the importance of kinetics in TCR signaling. Immunity 1998, 9, 817–826. doi:10.1016/s1074-7613(00)80647-0.

10. McKeithan, T.W. Kinetic proofreading in T-cell receptor signal transduction. Proc. Natl. Acad. Sci. USA 1995, 92, 5042–5046. doi:10.1073/pnas.92.11.5042.

11. Chakraborty, A.K.; Weiss, A. Insights into the initiation of TCR signaling. Nat. Immunol. 2014, 15, 798–807. doi:10.1038/ni.2940.

12. Siller-Farfan, J.A.; Dushek, O. Molecular mechanisms of T cell sensitivity to antigen. Immunol. Rev. 2018, 285, 194–205. doi:10.1111/imr.12690.

13. Courtney, A.H.; Lo, W.L.; Weiss, A. TCR signaling: Mechanisms of initiation and propagation. Trends Biochem. Sci. 2018, 43, 108–123. doi:10.1016/j.tibs.2017.11.008.

14. Wülfing, C.; Sumen, C.; Sjaastad, M.D.; Wu, L.C.; Dustin, M.L.; Davis, M.M. Costimulation and endogenous MHC ligands contribute to T cell recognition. Nature Immunology 2001, 3, 42–47. doi:10.1038/ni741.

15. Stefanová, I.; Dorfman, J.R.; Germain, R.N. Self-recognition promotes the foreign antigen sensitivity of naive T lymphocytes. Nature 2002, 420, 429–234. doi:10.1038/nature01146.

16. Dustin, M.L.; Cooper, J.A. The immunological synapse and the actin cytoskeleton: molecular hardware for T cell signaling. Nat. Immunol. 2000, 1, 23–29.

17. Cai, E.; Marchuk, K.; Beemiller, P.; Beppler, C.; Rubashkin, M.G.; Weaver, V.M.; Gerard, A.; Liu, T.L.; Chen, B.C.; Betzig, E.; Bartumeus, F.; Krummel, M.F. Visualizing dynamic microvillar search and stabilization during ligand detection by T cells. Science 2017, 356, eaal3118. doi:10.1126/science.aal3118.

18. Grakoui, A.; Bromley, S.K.; Sumen, C.; Davis, M.M.; Shaw, A.S.; Allen, P.M.; Dustin, M.L. The immunological synapse: a molecular machine controlling T cell activation. Science 1999, 285, 221–227.

19. Huppa, J.B.; Axmann, M.; Mörtelmaier, M.A.; Lillemeier, B.F.; Newell, E.W.; Brameshuber, M.; Klein, L.O.; Schütz, G.J.; Davis, M.M. TCR-peptide-MHC interactions in situ show accelerated kinetics and increased affinity. Nature 2010, 463, 963–967.

20. Weikl, T.R. Membrane-Mediated Cooperativity of Proteins. Annu. Rev. Phys. Chem. 2018, 69, 521–539.

21. Lipowsky, R. Lines of renormalization group fixed points for fluid and crystalline membranes. Europhys. Lett. 1988, 7, 255–261.

22. Nagle, J.F. Introductory Lecture: Basic quantities in model biomembranes. Faraday Discuss. 2013, 161, 11–29.

23. Dimova, R. Recent developments in the field of bending rigidity measurements on membranes. Adv. Colloid Interface Sci. 2014, 208, 225–234.

24. Liu, B.; Chen, W.; Evavold, B.D.; Zhu, C. Accumulation of dynamic catch bonds between TCR and agonist peptide-MHC triggers T cell signaling. Cell 2014, 157, 357–368. doi:10.1016/j.cell.2014.02.053.

25. Klein, L.; Kyewski, B.; Allen, P.M.; Hogquist, K.A. Positive and negative selection of the T cell repertoire: what thymocytes see (and don’t see). Nat. Rev. Immunol. 2014, 14, 377–391. doi:10.1038/nri3667.

26. Davis, S.J.; van der Merwe, P.A. The kinetic-segregation model: TCR triggering and beyond. Nat. Immunol. 2006, 7, 803–809. doi:10.1038/ni1369.

27. Choudhuri, K.; van der Merwe, P.A. Molecular mechanisms involved in T cell receptor triggering. Semin. Immunol. 2007, 19, 255–261. doi:10.1016/j.smim.2007.04.005.

28. Chang, V.T.; Fernandes, R.A.; Ganzinger, K.A.; Lee, S.F.; Siebold, C.; McColl, J.; Jönsson, P.; Palayret, M.; Harlos, K.; Coles, C.H.; Jones, E.Y.; Lui, Y.; Huang, E.; Gilbert, R.J.C.; Klenerman, D.; Aricescu, A.R.; Davis, S.J. Initiation of T cell signaling by CD45 segregation at ‘close contacts’. Nat. Immunol. 2016, 17, 574–582. doi:10.1038/ni.3392.

29. Pettmann, J.; Santos, A.M.; Dushek, O.; Davis, S.J. Membrane Ultrastructure and T Cell Activation. Front. Immunol. 2018, 9, 2152. doi:10.3389/fimmu.2018.02152.

30. Valitutti, S. The serial engagement model 17 years after: from TCR triggering to immunotherapy. Front. Immunol. 2012, 3, 272. doi:10.3389/fimmu.2012.00272.

31. Valitutti, S.; Müller, S.; Cella, M.; Padovan, E.; Lanzavecchia, A. Serial triggering of many T-cell receptors by a few peptide-MHC complexes. Nature 1995, 375, 148–151. doi:10.1038/375148a0.

32. Xu, G.K.; Hu, J.; Lipowsky, R.; Weikl, T.R. Binding constants of membrane-anchored receptors and ligands: A general theory corroborated Monte Carlo simulations. J. Chem. Phys. 2015, 143, 243136.

33. Krobath, H.; Rozycki, B.; Lipowsky, R.; Weikl, T.R. Binding cooperativity of membrane adhesion receptors. Soft Matter 2009, 5, 3354–3361.

34. Springer, T.A. Adhesion receptors of the immune system. Nature 1990, 346, 425–434.

35. Schmid, E.M.; Bakalar, M.H.; Choudhuri, K.; Weichsel, J.; Ann, H.S.; Geissler, P.L.; Dustin, M.L.; Fletcher, D.A. Size-dependent protein segregation at membrane interfaces. Nat. Phys. 2016, 12, 704.

36. Taylor, M.J.; Husain, K.; Gartner, Z.J.; Mayor, S.; Vale, R.D. A DNA-Based T Cell Receptor Reveals a Role for Receptor Clustering in Ligand Discrimination. Cell 2017, 169, 108–119.e20.

37. Chung, M.; Koo, B.J.; Boxer, S.G. Formation and analysis of topographical domains between lipid membranes tethered by DNA hybrids of different lengths. Faraday Discuss. 2013, 161, 333–345.

38. Choudhuri, K.; Wiseman, D.; Brown, M.H.; Gould, K.; van der Merwe, P.A. T-cell receptor triggering is critically dependent on the dimensions of its peptide-MHC ligand. Nature 2005, 436, 578–582. doi:10.1038/nature03843.

39. Milstein, O.; Tseng, S.Y.; Starr, T.; Llodra, J.; Nans, A.; Liu, M.; Wild, M.K.; van der Merwe, P.A.; Stokes, D.L.; Reisner, Y.; Dustin, M.L. Nanoscale increases in CD2-CD48-mediated intermembrane spacing decrease adhesion and reorganize the immunological synapse. J Biol Chem 2008, 283, 34414–34422.

40. Rossboth, B.; Arnold, A.M.; Ta, H.; Platzer, R.; Kellner, F.; Huppa, J.B.; Brameshuber, M.; Baumgart, F.; Schütz, G.J. TCRs are randomly distributed on the plasma membrane of resting antigen-experienced T cells. Nat Immunol 2018, 19, 821–827. doi:10.1038/s41590-018-0162-7.

41. Campi, G.; Varma, R.; Dustin, M. Actin and agonist MHC-peptide complex-dependent T cell receptor microclusters as scaffolds for signaling. J. Exp. Med. 2005, 202, 1031–1036.

42. Yokosuka, T.; Sakata-Sogawa, K.; Kobayashi, W.; Hiroshima, M.; Hashimoto-Tane, A.; Tokunaga, M.; Dustin, M.; Saito, T. Newly generated T cell receptor microclusters initiate and sustain T cell activation by recruitment of Zap70 and SLP-76. Nat. Immunol. 2005, 6, 1253–1262.

43. Mossman, K.D.; Campi, G.; Groves, J.T.; Dustin, M.L. Altered TCR signaling from geometrically repatterned immunological synapses. Science 2005, 310, 1191–1193.

44. Choudhuri, K.; Dustin, M.L. Signaling microdomains in T cells. FEBS Lett. 2010, 584, 4823–4831.

45. Sherman, E.; Barr, V.; Manley, S.; Patterson, G.; Balagopalan, L.; Akpan, I.; Regan, C.K.; Merrill, R.K.; Sommers, C.L.; Lippincott-Schwartz, J.; Samelson, L.E. Functional nanoscale organization of signaling molecules downstream of the T cell antigen receptor. Immunity 2011, 35, 705–720. doi:https://doi.org/10.1016/j.immuni.2011.10.004.

46. Pageon, S.V.; Tabarin, T.; Yamamoto, Y.; Ma, Y.; Nicovich, P.R.; Bridgeman, J.S.; Cohnen, A.; Benzing, C.; Gao, Y.; Crowther, M.D.; Tungatt, K.; Dolton, G.; Sewell, A.K.; Price, D.A.; Acuto, O.; Parton, R.G.; Gooding, J.J.; Rossy, J.; Rossjohn, J.; Gaus, K. Functional role of T-cell receptor nanoclusters in signal initiation and antigen discrimination. Proc. Natl. Acad. Sci. USA 2016, 113, E5454–63. doi:10.1073/pnas.1607436113.

47. Asfaw, M.; Rozycki, B.; Lipowsky, R.; Weikl, T.R. Membrane adhesion via competing receptor/ligand bonds. Europhys. Lett. 2006, 76, 703–709.

48. Różycki, B.; Lipowsky, R.; Weikl, T.R. Segregation of receptor–ligand complexes in cell adhesion zones: phase diagrams and the role of thermal membrane roughness. New J. Phys. 2010, 12, 095003.

49. Bachmann, M.F.; Barner, M.; Kopf, M. CD2 sets quantitative thresholds in T cell activation. J. Exp. Med. 1999, 190, 1383–1392. doi:10.1084/jem.190.10.1383.

50. Demetriou, P.; Abu-Shah, E.; McCuaig, S.; Mayya, V.; Valvo, S.; Korobchevskaya, K.; Friedrich, M.; Mann, E.; Lee, L.Y.; Starkey, T.; Kutuzov, M.A.; Afrose, J.; Siokis, A.; Meyer-Hermann, M.; Depoil, D.; Dustin, M.L. CD2 expression acts as a quantitative check-point for immunological synapse structure and T-cell activation. bioRxiv 2019, [https://www.biorxiv.org/content/early/2019/03/29/58944 doi:10.1101/589440.

51. Göhring, J.; Kellner, F.; Schrangl, L.; René Platzer, R.; Klotzsch, E.; Stockinger, H.; Huppa, J.B.; Gerhard J. Schütz, G.J. Temporal analysis of T-cell receptor-imposed forces via quantitative single molecule FRET measurements. bioRxiv 2020, https://doi.org/10.1101/2020.04.03.024299.

52. Schütz, G.; Huppa, J. How drag sharpens a T cell’s view on antigen. Proc Natl Acad Sci USA 2019, 116, 16669–16671.

53. Klotzsch, E.; Schuetz, G.J. Improved ligand discrimination by force-induced unbinding of the T cell receptor from peptide-MHC. Biophys. J. 2013, 104, 1670–1675. doi:10.1016/j.bpj.2013.03.023.

54. Hong, J.; Ge, C.; Jothikumar, P.; Yuan, Z.; Liu, B.; Bai, K.; Li, K.; Rittase, W.; Shinzawa, M.; Zhang, Y.; Palin, A.; Love, P.; Yu, X.; Salaita, K.; Evavold, B.D.; Singer, A.; Zhu, C. A TCR mechanotransduction signaling loop induces negative selection in the thymus. Nat. Immunol. 2018, 19, 1379–1390. doi:10.1038/s41590-018-0259-z.

55. Sibener, L.V.; Fernandes, R.A.; Kolawole, E.M.; Carbone, C.B.; Liu, F.; McAffee, D.; Birnbaum, M.E.; Yang, X.; Su, L.F.; Yu, W.; Dong, S.; Gee, M.H.; Jude, K.M.; Davis, M.M.; Groves, J.T.; Goddard, 3rd, W.A.; Heath, J.R.; Evavold, B.D.; Vale, R.D.; Garcia, K.C. Isolation of a structural mechanism for uncoupling T cell receptor signaling from peptide-MHC binding. Cell 2018, 174, 672–687.e27. doi:10.1016/j.cell.2018.06.017.

56. Limozin, L.; Bridge, M.; Bongrand, P.; Dushek, O.; van+der+Merwe, P.; Robert, P. TCR–pMHC kinetics under force in a cell-free system show no intrinsic catch bond, but a minimal encounter duration before binding. Proc. Natl. Acad. Sci. USA 2019, 116, 16943–16948.

57. Weikl, T.R.; Hu, J.; Kav, B.; Różycki, B. Binding and segregation of proteins in membrane adhesion: theory, modeling, and simulations. Advances in Biomembranes and Lipid Self-Assembly 2019, 30, 159–194. doi:https://doi.org/10.1016/bs.abl.2019.10.004.

58. Steinkühler, J.; Rozycki, B.; Alvey, C.; Lipowsky, R.; Weikl, T.R.; Dimova, R.; Discher, D.E. Membrane fluctuations and acidosis regulate cooperative binding of ‘marker of self’ protein CD47 with the macrophage checkpoint receptor SIRP*α*. J. Cell. Sci. 2019, 132, jcs216770.

59. Weikl, T.R.; Groves, J.T.; Lipowsky, R. Pattern formation during adhesion of multicomponent membranes. Europhys. Lett. 2002, 59, 916–922.

60. Weikl, T.R.; Lipowsky, R. Pattern formation during T-cell adhesion. Biophys. J. 2004, 87, 3665–3678.

61. Tsourkas, P.K.; Baumgarth, N.; Simon, S.I.; Raychaudhuri, S. Mechanisms of B-cell synapse formation predicted by Monte Carlo simulation. Biophys. J. 2007, 92, 4196–4208.

62. Knezevic, M.; Jiang, H.; Wang, S. Active Tuning of Synaptic Patterns Enhances Immune Discrimination. Phys. Rev. Lett. 2018, 121, 238101.

63. Qi, S.Y.; Groves, J.T.; Chakraborty, A.K. Synaptic pattern formation during cellular recognition. Proc. Natl. Acad. Sci. USA 2001, 98, 6548–6553.

64. Burroughs, N.J.; Wülfing, C. Differential segregation in a cell-cell contact interface: the dynamics of the immunological synapse. Biophys. J. 2002, 83, 1784–1796.

65. Raychaudhuri, S.; Chakraborty, A.K.; Kardar, M. Effective membrane model of the immunological synapse. Phys. Rev. Lett. 2003, 91, 208101.

66. Coombs, D.; Dembo, M.; Wofsy, C.; Goldstein, B. Equilibrium thermodynamics of cell-cell adhesion mediated by multiple ligand-receptor pairs. Biophys. J. 2004, 86, 1408–1423.

